# Impaired KDM2B-mediated PRC1 recruitment to chromatin causes neural stem cell senescence and ASD/ID-like behavioral deficits

**DOI:** 10.1101/2021.09.13.459918

**Authors:** Yuen Gao, Natalia Duque-Wilckens, Mohammad B Aljazi, Adam J Moeser, George I Mias, Alfred J Robison, Yi Zhang, Jin He

**Author notes:** To whom correspondence should be addressed: Yi Zhang, Jin He.

## Abstract

Autism spectrum disorder (ASD) and intellectual disability (ID) are neurodevelopmental diseases associated with various genetic mutations. Recent clinical studies report that chromosomal 12q24.31 microdeletions are associated with human ASD/ID. However, the causality and underlying mechanisms linking 12q24.31 microdeletions to ASD/ID pathogenesis remain undetermined. Here we show *Kdm2b*, one of the genes located in chromosomal 12q24.31, plays a critical role in maintaining neural stem cells (NSCs) in the developing mouse brain. Loss of the CxxC-ZF domain of KDM2B impairs its function in recruiting Polycomb repressive complex 1 (PRC1) to chromatin, resulting in de-repression of genes involved in cell apoptosis, cell cycle arrest, NSC premature senescence, and leading to the loss of NSC populations in the brain. Importantly, the *Kdm2b* mutation is sufficient to induce ASD/ID-like social and memory deficits in adult mice. Thus, our study reveals a critical role of an epigenetic factor KDM2B in normal brain development, a causality between the *Kdm2b* mutation and genesis of ASD/ID-like phenotypes in mice, and potential molecular mechanisms linking the function of KDM2B-PRC1 in transcriptional regulation and NSC senescence to the12q24.31 microdeletion-associated ASD/ID.

## Introduction

Autism spectrum disorder (ASD) is one of the most prevalent neurodevelopmental diseases (NDDs) associated with various genetic mutations^1^. Recent genetic studies have revealed that human ASD risk genes are enriched with genes encoding chromatin modifying factors and transcription regulators^2-5^, highlighting the importance of epigenetic factors and their function in regulating gene expression during normal brain development. However, the molecular and cellular mechanisms underlying the mutations of specific epigenetic factors and NDD genesis are largely unknown.

Recent genetic and clinical studies report that chromosomal 12q24.31 microdeletions are associated with NDDs in human patients^6-10^. Clinically, the patients display intellectual disability (ID), autistic behaviors, epilepsy, craniofacial deformity, and developmental delay, suggesting the genes located in 12q24.31 potentially have important functions for normal brain development. Further genetic mapping analysis identified a 445-kb common region in the chromosomal 12q24.31 microdeletion variants, in which one of the genes, *KDM2B*, was found to have reduced expression in patients compared to their healthy siblings, suggesting *KDM2B* was a candidate ASD pathogenic gene, and its loss-of-function mutations potentially contributed to the chromosomal 12q24.31 microdeletion-associated ASD/ID^8^.

KDM2B is a histone H3 lysine 36 di-methylation (H3K36me2)-specific demethylase^11^. In addition to its enzymatic activity, KDM2B interacts with Polycomb repressive complex 1 (PRC1) and recruits the KDM2B-PRC1 complex to chromatin through its CxxC zinc finger (CxxC-ZF) domain-mediated DNA binding to un-methylated CpG islands (CGIs) and CGI-associated gene promoters^12-14^. The recruitment of PRC1 to chromatin is essential for PRC1 to ubiquitylate histone H2A at gene promoters and repress gene expression in mammalian cells^15,16^. Although the mechanisms underlying the KDM2B-PRC1 complex-mediated transcriptional regulation have been well studied, the molecular and cellular mechanisms linking the *KDM2B* loss in 12q24.31 microdeletions to ASD/ID pathogenesis remain unknown.

To investigate the function of KDM2B in normal brain development and ASD/ID genesis, we generated a *Kdm2b* conditional mutant mouse line in which a mutant KDM2B with loss of its CxxC-ZF domain was introduced in the developing mouse brain. The genetic analysis demonstrated that the heterozygous *Kdm2b* mutation in the developing mouse brain impaired self-renewal of neural stem cells (NSCs), induced NSC senescence, and reduced NSC populations *in vitro* and *in vivo*, which was accompanied by increased cell apoptosis, cell cycle arrest, and precocious NSC differentiation. Importantly, behavioral studies showed that the heterozygous adult *Kdm2b* mutant mice displayed both ID-like memory deficits and core autistic-like behaviors, suggesting the *KDM2B* mutation-induced NSC loss in developing brains was likely to be a causative factor leading to the pathogenesis of 12q24.31 microdeletion-associated ASD/ID. At the molecular level, the *Kdm2b* mutation in NSCs resulted in decreased occupancy of KDM2B-PRC1 at gene promoters, and derepression of multiple genes involved in cell apoptosis, cell proliferation blockade, and promoting NSC differentiation. Thus, our study reveals a critical role of KDM2B in maintaining NSCs in the developing mouse brain and provides the molecular and cellular mechanisms linking the KDM2B-PRC1 function in transcriptional regulation to NSC senescence and 12q24.31 microdeletion-associated ASD/ID.

## Results

### Loss of KDM2B CxxC-ZF domain induces NSC senescence

Previous studies have shown that the CxxC-ZF domain of KDM2B is required for recruiting PRC1 to chromatin through its binding to non-methylated CpG-rich DNA sequences^12-14^. To examine the function of KDM2B-PRC1 *in vivo*, we generated a *Kdm2b* conditional mutant (*Kdm2b*-cMut) mouse line by inserting two *LoxP* elements into the exon 13-flanking sites at the *Kdm2b* gene locus (*Kdm2b*-cMut: *Kdm2b*^+/2f^). A CRE recombinase-mediated deletion of exon 13 resulted in altered splicing of mRNA encoding a mutant KDM2B protein with deletion of amino acids 552-626 encompassing the entire CxxC-ZF domain (Fig. S1a). The mutant KDM2B protein contained all functional domains except its CxxC-ZF domain, and thus lost its function in recruiting PRC1 to chromatin. To preclude mouse strain-specific effects on animal phenotypes, we backcrossed the wild-type *Kdm2b*^*+/2f*^ founders with C57BL/6 mice for more than five generations to reach a pure genetic background.

To examine the function of *Kdm2b* in central nervous system development, we crossed the *Kdm2b-*cMut mice with a neural progenitor cell (NPC)-specific Cre (Nestin-Cre) mouse line to generate neural *Kdm2b* conditional mutant mice (*Kdm2b*-Nes-cMut: *Kdm2b*^2f/+^;*Nestin-Cre*^*+/-*^). The following mating (*Kdm2b*^2f/2f^ × *Kdm2b*^2f/+^;*Nestin-Cre*^*+/-*^) only produced wild-type (*Kdm2b-*Nes-WT: *Kdm2b*^2f/2f^;*Nestin-Cre*^*-/-*^ or *Kdm2b*^2f/+^;*Nestin-Cre*^*-/-*^) and heterozygous (*Kdm2b-*Nes-Het: *Kdm2b*^2f/+^;*Nestin-Cre*^*+/-*^) mutant pups (Fig. S1b), suggesting the homozygous *Kdm2b* (*Kdm2b*^2f/2f^;*Nestin-Cre*^*+/-*^) mutation in the developing mouse brain was embryonic lethal. The *Kdm2b-*Nes-Het mice did not display any gross abnormalities and had similar postnatal growth to that of their wild-type littermates.

To investigate the function of KDM2B and its CxxC-ZF domain-mediated PRC1 recruitment in neural stem cells (NSCs) in the developing mouse brain, we isolated NSCs from the sub-ventricular zone (SVZ) at embryonic day 16 (E16) and examined their self-renewal by serial neurosphere formation assays. Compared to the wild-type NSCs, the *Kdm2b-*Nes-Het mutant NSCs formed primary neurospheres with smaller sizes in the first round of plating. After splitting into single cells, the wild-type NSCs formed secondary spheres with comparable sizes to the primary spheres, indicating the wild-type NSCs could maintain their self-renewing capacity under experimental conditions. In contrast, the *Kdm2b-*mutant cells failed to form secondary spheres in the second round of plating (Fig. 1a, b), suggesting the mutant NSCs were unable to maintain the stem cell property of self-renewal. Further senescence-associated beta-galactosidase (β-gal) staining assays showed that compared to the wild-type NSCs, the *Kdm2b-*Nes-Het mutant NSCs displayed a high percentage of β-gal+ cells at early passages, indicating the *Kdm2b* mutation induced premature NSC senescence (Fig. 1c, d). Taken together, these results suggested that heterozygous *Kdm2b* mutation with loss of its CxxC-ZF domain impaired NSC self-renewal by inducing premature cellular senescence *in vitro*.

**Figure 1.**
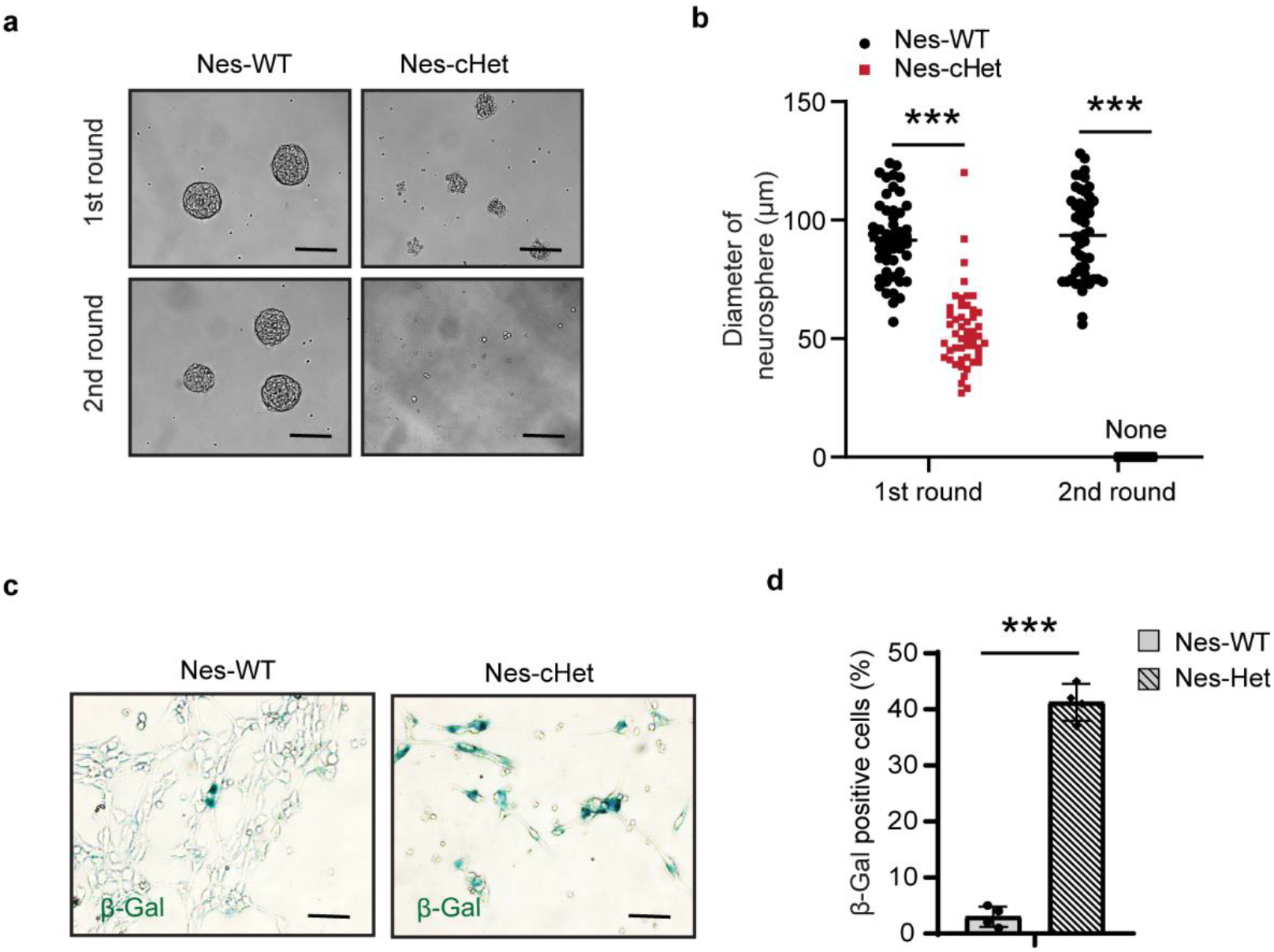
Loss of KDM2B CxxC-ZF domain induces NSC senescence. **a** Representative photos showing the size of primary and secondary neurospheres formed by wild-type and Kdm2b mutant NSCs in suspension culture; bar = 100 µm. **b** Dot plot showing the diameters of primary and secondary neurospheres formed by wild-type and Kdm2b mutant NSCs in suspension culture. *p-*values calculated using a two-way ANOVA; error bars represent mean ± SEM; \*\*\**p* < 0.001. **c** Representative photos showing the senescence-associated galactosidase-positive wild-type and *Kdm2b* mutant NSCs in monolayer culture; bar = 50 µm. **d** Bar plot showing the percentage of senescence-associated galactosidase-positive cells in each field. *p-*values calculated using a two-tailed *t*-test; error bars represent mean ± SEM; \*\*\**p* < 0.001.

### Loss of KDM2B CxxC-ZF domain induces cellular changes leading to impaired NSC self-renewal

Next, we examined the cellular changes that potentially could cause the NSC self-renewing defect observed in the *Kdm2b-*Nes-Het mutant cells, including cell proliferation, apoptosis, and precocious NSC differentiation. Bromodeoxyuridine (BrdU) incorporation assays showed that *Kdm2b-*mutant NSCs had significantly lower BrdU+ populations compared to wild-type cells, indicating the KDM2B mutation induced cell cycle arrest and blocked NSC proliferation (Fig. 2a, b). Moreover, terminal deoxynucleotidyl transferase dUTP nick end label (TUNEL) assays showed that the *Kdm2b-*mutant NSCs exhibited higher rate of apoptosis compared to the wild-type NSCs (Fig. 2c, d), suggesting that loss of CxxC-ZF domain of KDM2B induced cell apoptosis. To examine whether the *Kdm2b* mutation caused abnormal NSC differentiation, we stained the primary NSCs maintained in the NSC culture medium for the early differentiation marker GFAP. The results showed that the *Kdm2b-*Nes-Het mutant cells had a higher percentage of GFAP+ cells compared to the wild-type cells (Fig. 2e, f), suggesting the KDM2B mutation induced precocious NSC differentiation *in vitro*.

**Figure 2.**
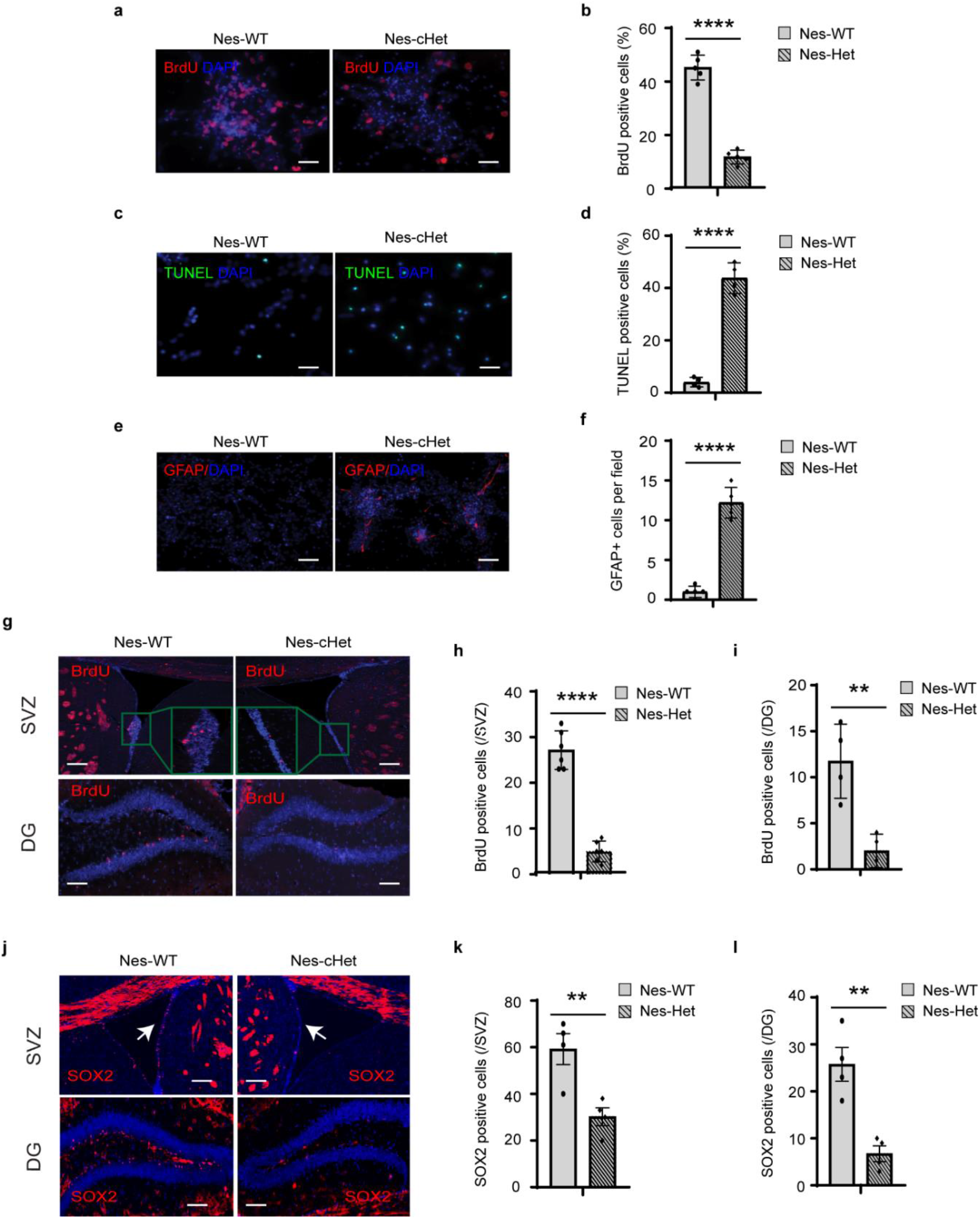
Loss of KDM2B CxxC-ZF domain induces cellular changes leading to NSC senescence. **a-b** Representative photos (a) and bar plots (b) showing the percentage of BrdU+ cells in the *Kdm2b*-Nes-WT and *Kdm2b*-Nes-Het NSCs cultured *in vitro*; bar = 20 µm. **c-d** Representative photos (c) and bar plots (d) showing the percentage of TUNEL+ apoptotic cells in the *Kdm2b*-Nes-WT and *Kdm2b*-Nes-Het NSCs cultured *in vitro*; bar = 20 µm. **e-f** Representative photos (e) and bar plots (f) showing the GFAP+ cells/field in the *Kdm2b*-Nes-WT and *Kdm2b*-Nes-Het NSCs cultured *in vitro;* bar = 20 µm. **g-i** Representative photos (g) and bar plots (h, i) showing the BrdU+ cells in the SVZ and DG of *Kdm2b*-Nes-WT and *Kdm2b*-Nes-Het mouse brains; bar = 100 µm. **j-l** Representative photos (j) and bar plots (k, l) showing the SOX2+ cells in the SVZ and DG of *Kdm2b*-Nes-WT and *Kdm2b*-Nes-Het mouse brains; bar = 100 µm. For panels b, d, f, h, i, k and l, 4–6 biological replicates per genotype were included in each experiment. *P-*values calculated using a two-tailed *t*-test; error bars represent mean ± SEM; \*\**p* < 0.01, \*\*\**p* < 0.001, **** *p* < 0.0001.

To examine whether the KDM2B mutation compromised NSC populations *in vivo*, we injected BrdU into adult mice and examined the BrdU-incorporated proliferating cells in brains. The results showed that compared to the wild-type littermates, the *Kdm2b-*Nes-Het mice had significantly fewer BrdU-incorporated cells in the SVZ and dentate gyrus (DG) of the hippocampus (Fig. 2g-i), which was consistent with lower numbers of SOX2+ NSCs in both regions (Fig. 2j-l). Grossly, although the mutant brain had normal morphology and similar weight to that of the wild-type, the hippocampus isolated from the mutant mice had ∼20% less weight than the wild-type littermates (Fig. S2a-c). Taken together, these results suggested the KDM2B mutation in the developing mouse brain induced cell apoptosis, repressed proliferation, and promoted precocious NSC differentiation, which subsequently led to the impaired NSC self-renewal and loss of NSC populations in the developing brain.

### *Kdm*2b mutation in the developing brain causes ASD/ID-like behavioral and memory deficits

Since the *Kdm2b* mutant mice had reduced NSC populations in the DG and smaller hippocampi, we further asked whether the KDM2B mutation in the developing mouse brain could lead to hippocampus-related memory deficits in adult mice. To this end, we performed the novel object recognition (NOR) test to examine object recognition memory. 24 hours after initial familiarization with two identical objects in an arena, the mice were allowed to explore the same arena in the presence of a familiar object and a novel object. The results showed that, compared to wild type mice, the *Kdm2b*-Nes-Het mice showed reduced discrimination index (t = 2.252, df = 28, *p* = 0.032) (Fig. 3a), indicating a deficit in long-term object recognition memory. The impaired memory was unlikely to be caused by hyperactivity or increased anxiety-like behaviors since the open field tests showed the mutant mice had comparable locomotor activity and exploring time in the center of the arena as their wild-type littermates (Fig. S3a, b). To further confirm the memory deficit observed in the mutant mice, we used the Morris water maze (MWM) to compare spatial memory between wild-type and *Kdm2b*-Nes-Het mice. Compared to the wild-type littermates, the mutant mice showed significantly delayed spatial learning during the first 5-day acquisition training phase (F1,188 = 8.067, *p* = 0.005), and they spent less time in the platform quadrant during the post-training probe tests (t= 2.732, df = 38, *p* = 0.0098) (Figs. 3b, d), suggesting the *Kdm2b*-Nes-Het mice had impaired spatial learning and memory.

**Figure 3.**
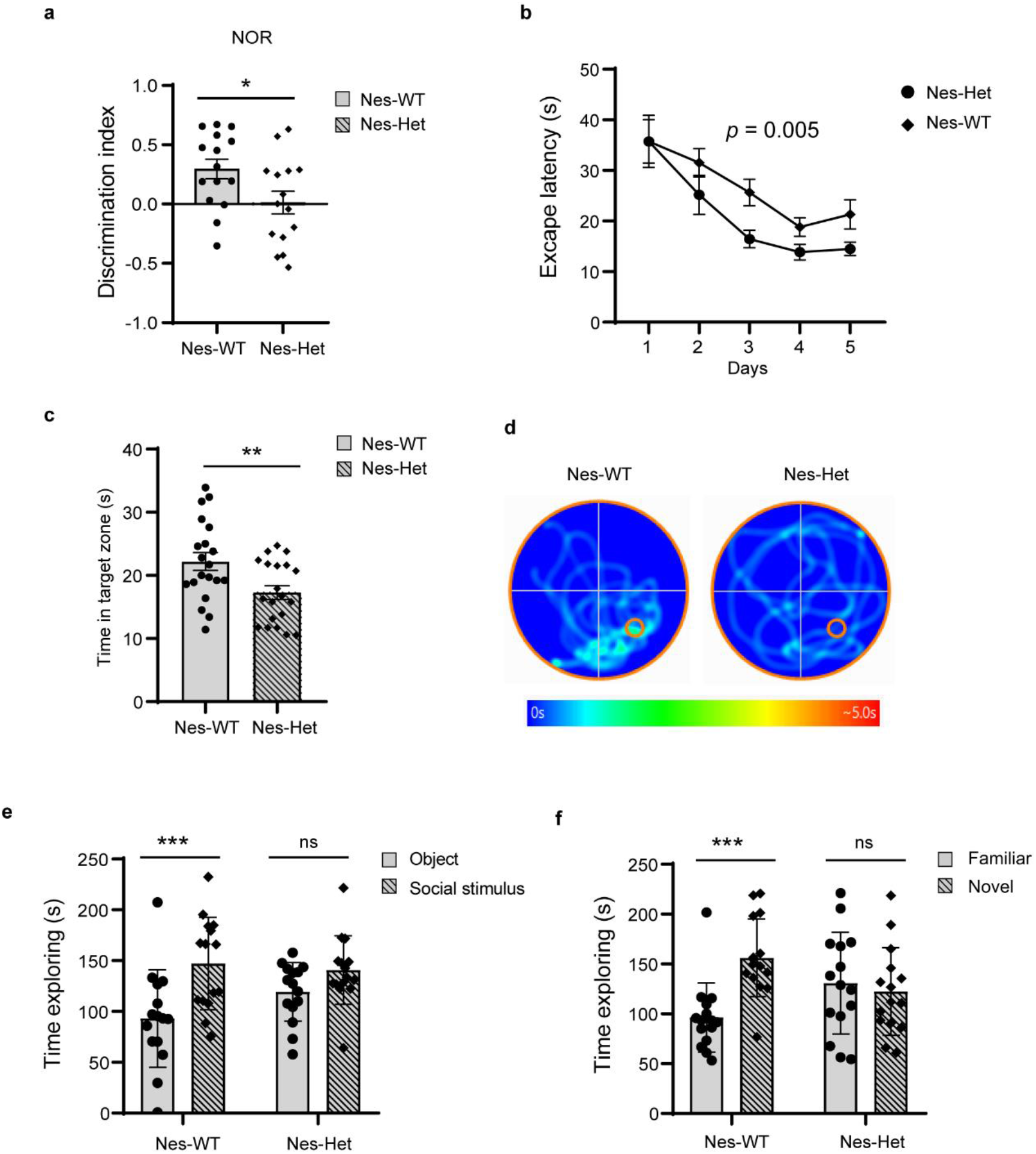
*Kdm*2b mutation in the developing brain causes ASD/ID-like behavioral and memory deficits. **a** Quantitative discrimination ratio of NOR tests. The discrimination ratio was calculated as: (time spent on the novel object-time spent on the familiar object)/total time. n = 15 for each genotype; *P-*values calculated using a two-tailed *t* test; error bars represent mean ± SEM; \**p* < 0.05. **b** The plot showing the escape latency during the 5-day training phase. n = 15 for each genotype; *P-*values calculated using two-way ANOVA; error bars represent mean ± SEM. **c** The time in the target zones during the post-training probe tests. n = 15 for each genotype; *P-*values calculated using a two-tailed *t* test; error bars represent mean ± SEM; \*\**p* < 0.01. **d** Representative heatmap showing the time in the target zone during the post-training probe test. The red circle represents the location of platform. **e** Quantitative results showing the time wild-type and *Kdm2b*-Nes-Het mice spent in the chamber containing a social partner in the 3-chamber sociability tests. N = 15 for each genotype; *P-*values calculated using two-way ANOVA; error bars represent mean ± SEM; \*\*\**p* < 0.001, ns = not significant. **f** Time spent in the chamber containing a novel animal in the 3-chamber social novelty tests. n = 15 for each genotype; *P-*values calculated using two-way ANOVA; error bars represent mean ± SEM; \*\*\**p* < 0.001, ns = not significant.

To examine whether the *Kdm2b* mutant mice developed autistic-like behaviors, we performed a three-chamber test to examine voluntary exploration of a social vs. a non-social stimulus (sociability) and the voluntary exploration of a familiar vs. a novel social stimulus (social novelty/social memory)^17,18^. In the sociability portion of the test, the results showed main effects of stimulus type (F1,56 = 13.48, *p* = 0.0005). Planned comparisons revealed that the wild-type controls spent more time with the social stimulus than with the object (t = 3.713, df = 56, *p* = 0.0009), representing normal sociability. In contrast, the *Kdm2b*-Nes-Het mice showed a reduced preference for the social stimulus, suggesting impaired sociability (Fig. 3e). In the social novelty portion of the test, there was an effect of genotype by stimulus interaction (F1,68 = 7.929, *p* = 0.006). Planned comparisons showed that the wild-type mice preferred a novel over a familiar animal (*p* = 0.0001) (Fig. 3f), indicating preference for social novelty. In contrast, the *Kdm2b*-Nes-Het mice showed no preference, indicating loss of social interest or reduced social memory.

Thus, the collective behavioral tests revealed that the *Kdm2b*-Nes-Het mice with loss of KDM2B CxxC-ZF domain in the developing mouse brain displayed ASD/ID-like behavioral deficits, including impaired cognitive and spatial memory, reduced sociability, and impaired social memory.

### Loss of KDM2B CxxC-ZF domain depresses the genes involved in impaired NSC self-renewal

Next, we set out to investigate the molecular mechanisms underlying the phenotypic changes in the *Kdm2b* mutant NSCs by examining transcriptome changes. Consistent with the function of KDM2B-PRC1 in transcriptional repression, RNA-seq analysis revealed that the *Kdm2b-*Nes-Het mutant NSCs had more upregulated (n = 302) than the downregulated (n = 191) genes (cutoff: RPKM ≥ 3, fold changes ≥ 2, FDR < 0.05) (Fig. S4a). The gene ontology (GO) enrichment analysis showed that the upregulated genes could be divided into three main functional groups (cutoff: FDR < 0.05): (i) inducing cell apoptosis; (ii) repressing cell proliferation; and (iii) inducing neural development (Fig. 4a, and Tables S1, S2), which was consistent with the increased apoptosis, reduced cell proliferation, and precocious NSC differentiation of *Kdm2b-*Nes-Het mutant NSCs (Fig. 2a-f). This suggested that the genes aberrantly upregulated in the *Kdm2b-*Nes-Het mutant NSCs were likely to play a major role in inducing the phenotypic changes observed in the *Kdm2b* mutant cells. In contrast, the GO enrichment analysis showed that the downregulated genes were involved in initiation of DNA replication (Fig. 4b), suggesting the decreased NSC proliferation could be also caused by reduced genes involved in mitotic division.

**Figure 4.**
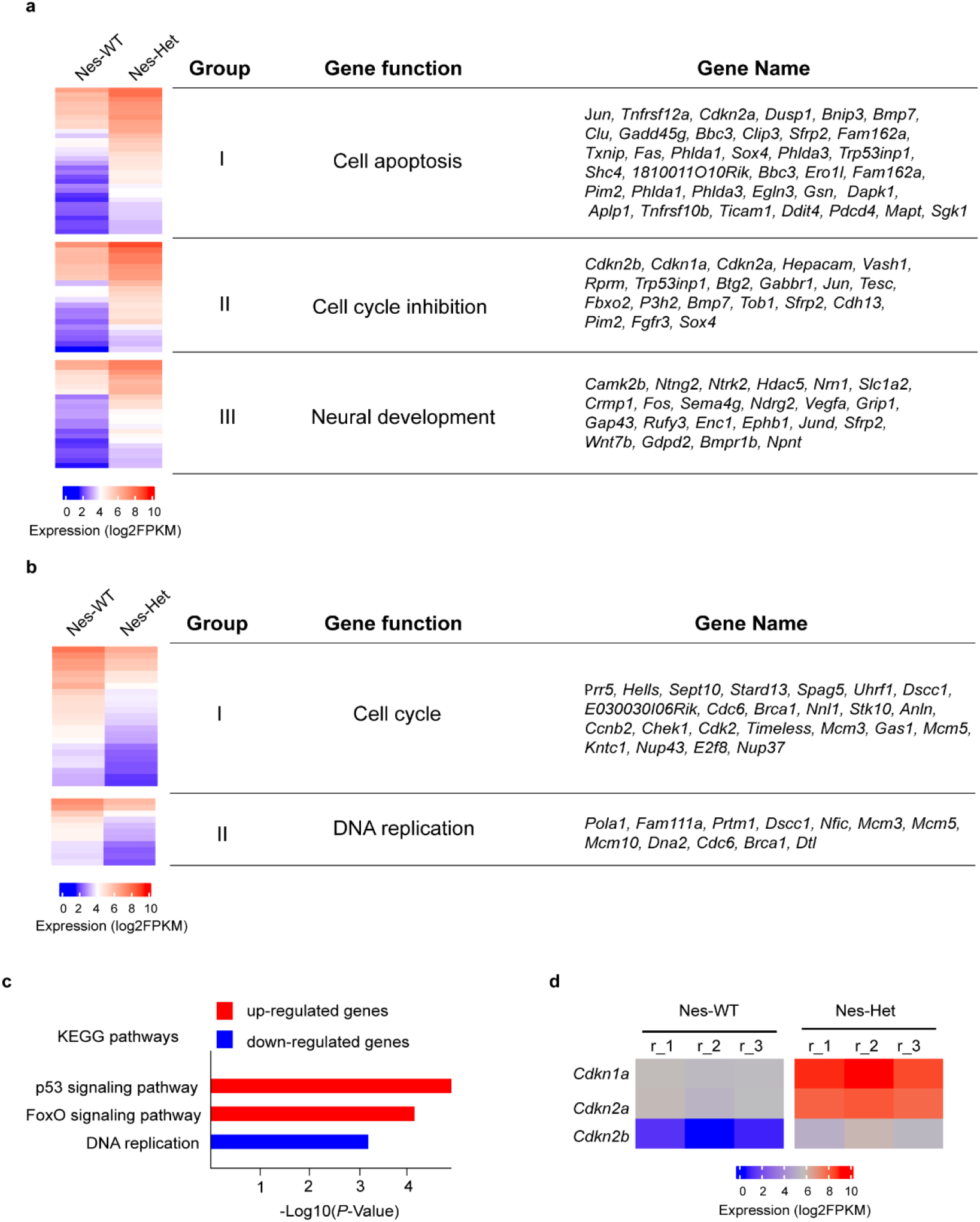
Loss of KDM2B CxxC-ZF domain depresses the genes involved in NSC senescence. **a** The up-regulated genes in the *Kdm2b*-Nes-cKO NSCs and their related gene functions. **b** The down-regulated genes in the *Kdm2b*-Nes-cKO NSCs and their related gene functions. **c** KEGG pathway enrichment analysis showing the functional pathways related to the up- and down-regulated genes in the *Kdm2b*-Nes-cKO NSCs during differentiation. **d** Heatmap showing the expression of *Cdkn1a, Cdkn2a*, and *Cdkn2b* in the *Kdm2b*-Nes-WT and *Kdm2b*-Nes-cKO NSCs.

Further, Kyoto Encyclopedia of Genes and Genomes (KEGG) pathway enrichment analysis showed the upregulated genes in the *Kdm2b-*Nes-Het mutant were involved in the activation of p53 and FOXO signaling pathways that induce both cell apoptosis and cell cycle arrest^19-21^, whereas the downregulated genes were involved in regulating cell cycle (Fig. 4c and Tables S3, S4). Notably, *Cdkn2a* and *Cdkn2b* in the *Ink4a-Arf* locus and *Cdkn1a*, which play a central role in inducing both RB-mediated cell cycle arrest and p53-mediated apoptosis^22,23^, were highly upregulated in the *Kdm2b-*Nes-Het mutant NSCs (Figs. 4d, S4b), consistent with previous findings that Polycomb-mediated epigenetic modification was critical for repressing the *Ink4a*-*Arf* locus to maintain NSCs in the developing mouse brain^24-27^.

Thus, the collective results suggested that at the molecular level, the KDM2B mutation with loss of its CxxC-ZF domain derepressed the genes involved in apoptosis, cell proliferation blockade, and precocious NSC differentiation, which led to impaired NSC self-renewal, NSC senescence, and loss of NSC populations in the developing brain.

### Loss of KDM2b CxxC-ZF domain impaired the PRC1 recruitment to chromatin

To examine whether the derepressed gene expression in the *Kdm2b-*Nes-Het mutant NSCs was caused by impaired KDM2B-mediated PRC1 recruitment to chromatin, we performed chromatin immunoprecipitation coupled with next-generation sequencing (ChIP-seq) assays to examine the occupancy of KDM2B and PRC1 core component RING1B in wild-type and *Kdm2b-*Nes-Het mutant NSCs. The results showed that genome-wide KDM2B occupancy at the 7,125 KDM2B binding sites was reduced to ∼ 50% in the *Kdm2b-*Nes-Het mutant NPCs compared to that in the wild-type cells, confirming that loss of the KDM2B CxxC-ZF domain impaired its chromatin binding. To examine whether the mutant KDM2B reduced the occupancy of the PRC1 core component RING1B at gene promoters, we compared the total RING1B occupancy at the promoters of differentially expressed gene in wild-type and *Kdm2b-*Nes-Het mutant NSCs (Fig. S4a). The results showed that RING1B occupancy was reduced for ∼25% at the promoters of upregulated genes in the *Kdm2b-*Nes-Het mutant NSCs, whereas its occupancy at downregulated gene promoters had a slight increase (Figs. 5b, S5). Specifically, both KDM2B and RING1B occupancy was reduced at the *Ink4a-Arf* locus (Fig. 5c), which was consistent with the upregulated expression of *Cdkn2a* and *Cdkn2b* in the *Kdm2b-*Nes-Het mutant NSCs (Figs. 4d, S4a). Taken together, these results suggested that aberrant gene expression in the *Kdm2b-*Nes-Het mutant NSCs was a direct outcome of reduced KDM2B-mediated PRC1 recruitment and impaired PRC1-mediated transcriptional repression.

**Figure 5.**
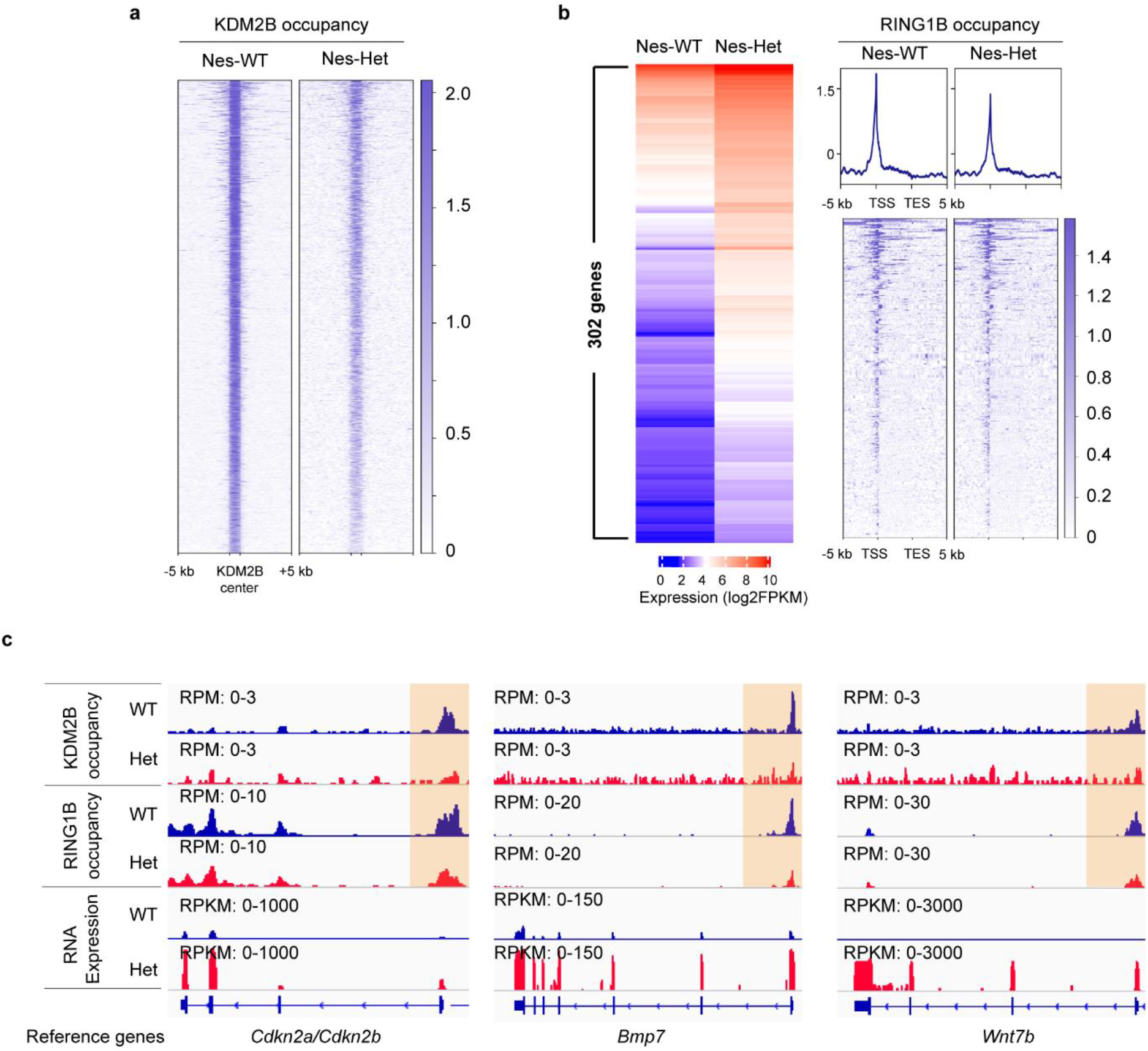
Loss of KDM2b CxxC-ZF domain impaired the PRC1 recruitment to chromatin. **a** Heatmap showing KDM2B occupancy at KDM2B binding sites and their 10-kb flanking sites in the *Kdm2b*-Nes-WT and *Kdm2b*-Nes-cKO NSCs. **b** Heatmaps and plots showing the 302 upregulated genes in the *Kdm2b*-Nes-cKO NSCs (left panel) and RING1B occupancy at the gene code regions and their 10kb flanking regions (right panel). TSS: transcriptional starting sites; TES: transcriptional ending sites. **c** IGV genome browser view of KDM2B occupancy, RING1B occupancy, and expression levels of *Cdkn2a/2b, Bmp7*, and *Wnt7b* genes.

## Discussion

In this study, we used a *Kdm2b* conditional mutant mouse model to investigate two questions related to chromosomal 12q24.31 microdeletion-associated ASD/ID: (i) Is the loss-of function mutation of KDM2B a causative driver leading to ASD/ID-like behavioral and memory deficits? (ii) What are the molecular mechanisms linking *Kdm2b* mutations to ASD/ID pathogenesis?

At the cellular level, the *Kdm2b* mutant NSCs displayed impaired self-renewal and underwent premature cellular senescence (Fig. 1), which was accompanied with increased cell apoptosis, reduced cell proliferation, and premature NSC differentiation. Correspondingly, the mutant adult mice had markedly reduced SOX2+ and BrdU+ proliferating NSCs in the SVZ and DG (Fig. 2), suggesting the CxxC-ZF domain of KDM2B is required for maintaining the stem cell populations in the developing mouse through preventing premature NSC senescence.

Previous studies have shown that the CxxC-ZF domain of KDM2B mediates the recruitment of PRC1 to chromatin through its binding to unmethylated CpG-rich DNA sequences^12-14^. After binding to chromatin, PRC1 catalyzes the mono-ubiquitination of histone H2A at gene promoters and mediates transcriptional silencing^16,28^. Consistent with these findings, transcriptome analyses showed that loss of the CxxC-ZF domain of KDM2B in NSCs derepressed genes involved in cell apoptosis, cell proliferation blockade, and NSC premature differentiation. Specifically, *Cdkn2a* and *Cdkn2b* in the *Ink4a-Arf* locus, which are located at the apex of signaling pathways to activate RB and p53-mediated cell cycle arrest and apoptosis^22,23^, were highly upregulated in the *Kdm2b* mutant NSCs (Fig. 4). Furthermore, ChIP-seq analysis showed that the occupancy of KDM2B and RIGN1B was largely reduced at the promoters of activated genes (Fig. 5), suggesting that de-repression of genes in the *Kdm2b* mutant NSCs was a direct outcome caused by reduced KDM2B-mediated PRC1 recruitment to chromatin. Taken together, these results suggest that the KDM2B mutation with loss of its CxxC-ZF domain impaired the KDM2B-mediated PRC1 recruitment to chromatin, compromised the function of PRC1 in transcriptional silencing, and led to de-repression of genes. The overall consequences of these molecular changes are the defective NSC self-renewal and premature senescence in the *Kdm2b* mutant brain.

Further behavioral studies showed that the KDM2B mutation with loss of its CxxC-ZF domain in the developing mouse brain was sufficient to cause ID-like cognitive and spatial memory deficits as well as autistic-like deficits in social behaviors (Figure 4), suggesting that loss-of-function mutation of KDM2B in the developing mouse brain was likely to be a causative factor leading to 12q24.31 microdeletion-associated ASD/ID pathogenesis. Although currently the roles of NSCs in ASD/ID pathogenesis remain undetermined, earlier studies have shown that loss of NSCs in young postnatal mice causes progressive memory impairment^29^, which is consistent with our findings that the KDM2B mutation-induced cellular senescence and loss of NSC populations in the developing brain result in ID-like cognitive and spatial memory deficits. Based on these results, we propose that KDM2B is critical for the maintenance of NSC populations in the developing mouse brain by recruiting PRC1 to chromatin to silence the genes involved in cell apoptosis, cell cycle blockade, and precocious NSC differentiation. The mutant KDM2B with loss of its CxxC-ZF domain causes aberrant gene expression, premature NSC senescence, and loss of NSC populations in the developing brain, which subsequently causes the ASD/ID-like behavioral and memory deficits in adult mice (Figure 6).

**Figure 6.**
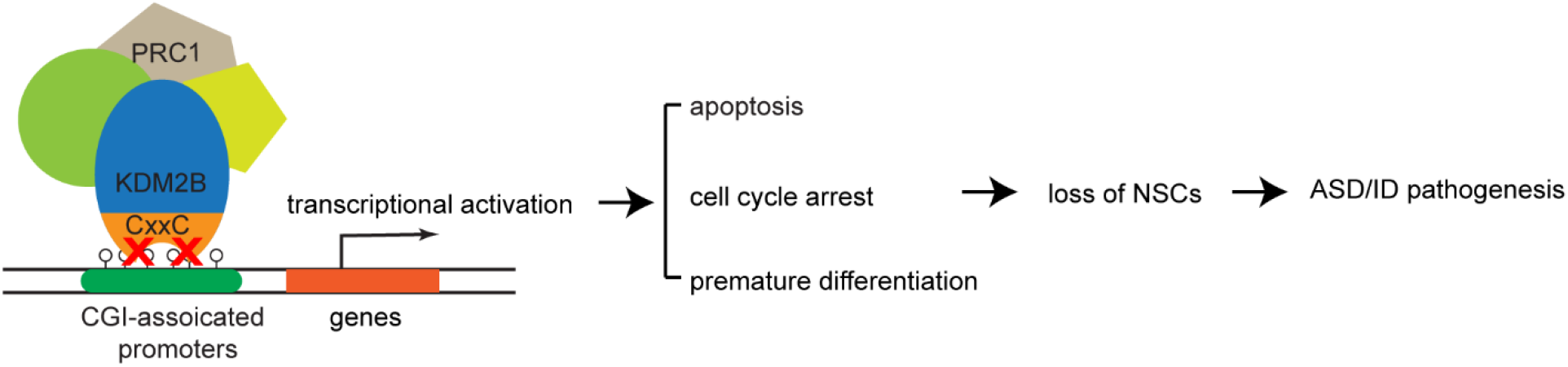
Proposed model for the loss-of-function of KDM2B-induced ASD/ID pathogenesis. KDM2B is critical for the maintenance of NSC populations in the developing mouse brain by recruiting PRC1 to chromatin to silence the genes involved in cell apoptosis, cell cycle blockade, and precocious NSC differentiation. The KDM2B mutation with loss of its CxxC-ZF domain derepresses genes involved in apoptosis, cell cycle arrest, and premature NSC differentiation, which leads to loss of NSCs and ASD/ID pathogenesis.

Our current study is exciting, but has some limitations: (i) unlike the loss of the entire protein in the chromosomal 12q24.31 microdeletions, the heterozygous *Kdm2b* mutant mice used in this study generate both wild-type and mutant KDM2B proteins in the brain, in which the mutant proteins might have a dominant-negative effect to exacerbate the phenotypes; (ii) although the *Kdm2b* mutation-induced NSC senescence and loss of NSC populations in the brain are likely to play a major role in causing the ASD/ID-like abnormalities in this mouse model, our current study does not exclude other mechanisms, such as aberrant neural lineage differentiation caused by de-repression of lineage-specific transcription factors, which might also be involved in the genesis of ASD/ID-like phenotypes.

Finally, our mouse model has shown that the impaired KDM2B-mediated recruitment of PRC1 to chromatin is sufficient to induce NSC senescence and deplete NSC populations in the brain, which provides a useful genetic tool to study the functions of NSCs in adult neurogenesis, brain aging, and brain degenerative diseases by introducing the KDM2B mutation to deplete adult NSC populations in the brain.

## Materials and Methods

### Mice

The *Kdm2b* conditional mutant mice were generated in Dr. Yi Zhang lab. The *Kdm2b* conditional mutant target construct was generated by modifying the BAC clone (RP23-214I6) based on the Recombneering method^30^. Two LoxP elements were inserted into the exon 13-flanking sites. The targeting construct was electroporated into the C57BL/6:129 hybrid murine ES cells. Homologous recombinant ES cell clones were identified by PCR-based genotyping and injected into blastocysts. The genetically modified ES cells were micro-injected to blastocysts and transferred to the uterus of CD-1 pseudo-pregnant females to generate chimeric founder mice. The chimeric founder mice were crossed to a FLP recombinase mouse line to remove the FRT-flanked selection cassette. All mice were backcrossed to C57BL/6 mice for at least five generations to reach a pure C57BL/6 background before further mating to specific Cre lineages. Mice were housed under standard conditions (12h light: 12h dark cycles) with food and water *ad libitum*. The data obtained from all embryos were pooled for analysis without discrimination of sexes. All mouse experiments were performed with the approval of the Michigan State University Institutional Animal Care & Use Committee.

### Mouse breeding strategy

All mice were backcrossed to C57BL/6 mice for at least five generations to reach a pure C57BL/6 background. (1) Generating *Kdm2b*-Nestin-cMut mice: The *Kdm2b* neural conditional mutant mice were generated by mating *Ash1L* floxed mice with Nestin-cre mice (B6.Cg-Tg (Nes-cre) 1Kln/J, The Jackson Laboratory). The wild-type (*Kdm2b*^2f/2f^;*Nestin-Cre*^*-/-*^) and heterozygous (*Kdm2b*^2f/+^;*Nestin-Cre*^*+/-*^) mice were generated by *Kdm2b*^2f/2f^;*Nestin-Cre*^*-/-*^ (female) × *Kdm2b*^2f/+^;*Nestin-Cre*^*+/-*^ (male) mating.

### Genotyping

Genomic DNA was extracted from mouse tails with a lysis buffer of 0.01M NaOH. After neutralizing with Tris-HCl (PH 7.6), the extracted genomic DNA was used for genotyping PCR assays. Primers used for genotyping are listed in Supplementary Table 5.

### Isolation, culture, and neurosphere formation of neural stem cells

The neural tissues were isolated from the subventricular zone of E16 brains and dissociated with 0.05% trypsin-EDTA (Life Technologies) at 37 °C for 15 min. The reaction was stopped by trypsin inhibitor (10 mg/ml, Worthington Biochemical Corporation). The dissociated cells were washed with cold PBS for 3 times and plated onto the non-coated petri dishes in the proliferation medium [Neurobasal medium (Life Technologies) supplemented with 1× B27 supplement (Gibco), 1x GlutaMAX (Life Technologies), 20 ng/ml murine epithelial growth factor (Peprotech), 20 ng/ml basic fibroblast growth factor (Peprotech), and 100U/ml penicillin/streptomycin (Life Technologies)]. For first round neurosphere formation assay, the neurospheres were formed by newly isolated primary culture (passage 0) neural progenitor cells after culturing for 7 days *in vitro*. The second round neurosphere formation assay was carried out by dissociating the first round neurospheres to single cells (passage 1) and culturing for another 7 days. The diameters of neurospheres were measured using a Zeiss Axio Imager microscope (Carl Zeiss GmbH, Oberkochen, Germany).

### NSC proliferation and differentiation assays

To study cell proliferation, we dissociated NSCs with trypsin and plated them on poly-L-ornithin/laminin-coated glass coverslips at a density of 50,000 cells/well in proliferation medium (see above). At 20 h post-plating, 5 μM 5-bromo-2’-deoxyuridine (BrdU, Sigma-Aldrich) was added into the culture medium for 10 h. NSCs were then washed with PBS and fixed with 4% paraformaldehyde for 30 min at room temperature, followed by immunohistochemical analysis. To detect BrdU incorporation, fixed cells were pretreated with 1.0 M HCl for 30 min at 37°C, and then washed with borate buffer (pH 8.5) for 30 min. To induce differentiation of neural progenitor cell, the cultured neurospheres were collected and re-plated onto the Poly-L-Ornithine (R&D systems) and Laminin (Corning)-coated coverslips to form a monolayer culture. The neural progenitor cells were passaged at a 1:5 ratio every 3 days. The NPC monolayer cells were gently washed with PBS 3 times and cultured under differentiation medium [Neurobasal medium supplemented with 1× N2 supplement (Giboco), 1x GlutaMAX (Life Technologies)]. Before immunocytochemistry staining, monolayer cultured cells were washed with DPBS and fixed with 4% PFA for 10 min. After pre-treatment, cells were pre-blocked with DPBS containing 5% normal donkey serum and 0.2% Triton X-100 for 30 min, followed by overnight incubation with primary antibody. The antibodies were rat anti-BrdU (1:500, MCA2060, Bio-Rad) or rabbit anti-GFAP (1:300, 12389, Cell Signaling technology). After washing with DPBS, cells were incubated with secondary antibodies R-Phycoerythrin AffiniPure F (ab’) 2 Fragment Donkey Anti-Rat IgG (1:300; 712-116-153; Jackson ImmunoResearch) or Alexa Fluor 568 donkey anti-rabbit IgG (1:300, Life Science technology) for 1 h and mounted using Vectorshield mounting media with DAPI (H1200, Vector Laboratories). Images were captured using a Zeiss Axio Imager microscope (Carl Zeiss GmbH, Oberkochen, Germany) and an AxioCam HRc camera (Carl Zeiss GmbH) with image acquisition via Zeiss Zen Pro software (v.2.3; Carl Zeiss GmbH).

### TUNEL assay

In Situ Cell Death Detection Kit (11684795910, Sigma-aldrich) was used to evaluate cell death in monolayer cultured neural progenitor cells according to the manufacturer’s instruction. Fluorescence images were captured by Zeiss Axio Imager microscope (Carl Zeiss GmbH, Oberkochen, Germany).

### Senescence-associated β-galactosidase staining assay

Senescence β-Galactosidase Staining Kit (9860, Cell Signaling technology) was applied to measure the senescent monolayer cultured neural progenitor cells according to the manufacturer’s instruction. Images were captured by Zeiss Axio Imager microscope (Carl Zeiss GmbH, Oberkochen, Germany).

### *In vivo* BrdU incorporation assay

4-5 week old mice were injected with 100 mg/kg BrdU (Sigma-Aldrich) intraperitoneally once every 3 days for 6 total doses. Seven days after the final injection, mice were anesthetized via ketamine (80 mg/kg) and xylazine (6 mg/kg) intraperitoneally, perfused intracardially with ice-cold solutions of 10 ml of DPBS followed by 10 ml of 4% PFA in PBS. Brains were extracted and fixed in 4% PFA for 24 h at 4°C, then embedded in paraffin. Coronal sections (5 μm) were harvested using a microtome.

### Immunohistology assays

Mouse tissues were fixed in 4% PFA in PBS overnight at 4 °C and embedded in paraffin. For immunofluorescence, tissue sections of 5 μm were cut, dewaxed, and rehydrated. Antigen retrieval was performed by microwaving the sections in 0.01 M sodium citrate buffer (pH 6.0) for 4 min. Tissue sections were blocked in 5% normal donkey serum (NDS) for 30 min after washing with PBS. Tissue sections were then incubated with rabbit anti-Sox2 (1:200, 3728, Cell Signaling technology) diluted in 5% NDS overnight at 4 °C. After washing with PBS, sections were incubated with Alexa Fluor 568 donkey anti-rabbit IgG (1:300, Life Science technology) for 1 h and mounted using Vectorshield mounting media with DAPI (H1200, Vector Laboratories). Alternately, when IHC was required for BrdU visualization, sections were sequentially subjected to HCl treatment, incubated in ice-cold 1 N HCl for 10 min, in 2 N HCl for 10 min at room temperature, and in 2 N HCl for 40 min at 37°C. After finishing the above steps, 1x PBS was used to wash the sections. Sections were then processed as described in neural progenitor cell proliferation assay.

### Behavioral tests

All behavioral tests were performed on littermate wild-type and heterozygous *Kdm2b*-Nes-cKO mice. The mice were labelled by ear-tags. All the behavioral tests were carried out by researchers blinded to the genotypes of animals during the behavioral tests.

### Open field test

The open-field apparatus consisted of a custom-made, square white polyvinylchloride foam box (38 × 38 × 35cm). Mouse behavior was recorded for the first 30 minutes of habituation to measure time spent in open field and distance traveled with a digital CCD camera connected to a computer running an automated video tracking software package (ANYMaze; Stoelting).

### Novel object recognition test (NOR)

NOR was assessed using a 3-day paradigm that included habituation, training, and testing as described previously^31-34^. Each day, mice were acclimated for 60 min to the behavioral testing room before assessment. All tests were performed under red lights, and behaviors were video recorded and automatically scored using Clever Sys. During habituation (day1), mice were placed into the open field apparatus, a square white polyvinylchloride foam 38 × 38 × 35 cm box, for 60 min while video recorded. For training (day 2), two identical objects consisting of miniature wheels, knobs, spark plugs, or Lego blocks were placed in opposite corners of the open field apparatus, and the animals were allowed to explore the objects for 60 min. The object pairs used were counterbalanced across treatments. For testing (day 3), mice were placed in the same apparatus, but this time one object of the pair was replaced with another dissimilar object (novel object), and mice were allowed to freely explore for 5 minutes. Their behavior was recorded, and the time the mice spent with their nose oriented towards the object within 3.5cm of the object edge was considered exploration time. Throughout testing, objects and apparatus were cleaned with 70% ethanol between trials. Discrimination index was calculated as:

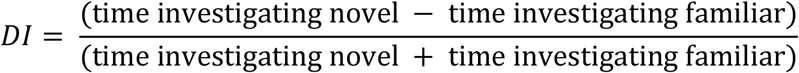

### Sociability and preference for social novelty test

This test was adapted from Crawley’s sociability and preference for social novelty protocol, which consists of three phases^17,18^. Mice were acclimated for 60 min to the behavioral testing room under red lights before testing. The behaviors during all three phases were video recorded and automatically scored using Clever Sys. In phase I (habituation), the experimental mouse was placed in the center of a three-chamber apparatus (polyvinylchloride, 60 × 40 × 22 cm) and allowed to freely explore for 5 minutes. During this time, the mouse had free access to all three chambers, which are connected by small openings at the bottom of the dividers. In phase 2 (sociability), two identical, wire cup-like containers were placed one in each of the side chambers. In this phase, an unfamiliar same-sex mouse was placed in one of the containers (“social stimulus”), while the other remained empty (“object”). The experimental mouse was allowed to freely explore the three chambers again for 5 minutes. In phase 3 (social memory), the container with the mouse (now “known”) was moved to the opposite chamber, and a new same-sex mouse (“unknown”) was placed in the other container. The experimental mouse was allowed to freely explore the three chambers for 5 minutes. Throughout testing, objects and apparatus were cleaned with 70% ethanol between trials. For analysis, the time with total body spent in each of the three chambers was recorded.

### Morris water maze

Animal activity was measured by a video-based tracking system (ANYMaze; Stoelting). The pool was filled with opaque water (by adding washable white paint) and surrounded by extra maze cues. During the visible platform training, the escape platform (10 cm in diameter) was placed in the center of a designated quadrant with its top positioned 1 cm above the water surface, and the platform was marked by a flag. Mice were trained by four trials per day for 2 d. The four trials were performed with 2 min interval between the trials [intertrial interval (ITI)]. Mice were allowed to navigate in the circular pool for up to 60 s until they found the platform. Mice were allowed to stay on the platform for 60 s. If mice failed to find and land on the platform within 60 s, they were manually guided to the platform. The visible platform was randomly placed in different locations for each trial. After the visible platform training, the mice were further trained by the hidden platform paradigm, during which the platform was placed 1 cm beneath the opaque water. Mice were trained by four trials (with 2 min ITI) per day for 5 d. For each trial, mice were dropped into the pool randomly from four different designated start points. The time each mouse spent to land on the platform was scored as escape latency. Probe trials were performed 1 d after the hidden platform training. With the escape platform removed, the mice were allowed to swim in the pool for 60 s. The time spent in target quadrant, number of crossings of the target quadrant, and the location of the hidden platform were recorded.

### RT-qPCR assays

RNA was extracted and purified from cells using QI shredder (Qiagen) and RNeasy (Qiagen) spin columns. Total RNA (1 µg) was subjected to reverse transcription using Iscript reverse transcription supermix (Bio-Rad). cDNA levels were assayed by real-time PCR using iTaq universal SYBR green supermix (Bio-Rad) and detected by CFX386 Touch Real-Time PCR detection system (Bio-Rad). Primer sequences for qPCR are listed in Supplementary Table 5.

### ChIP-Seq sample preparation

For KDM2B, RING1B, uH2A ChIP, primary wild-type and *Kdm2b*^1f/+^ NSCs were fixed with 2 mM ethylene glycol bis[succinimidylsuccinate] (Thermo Scientific) for 1 hour, followed by 10 min in 1% formaldehyde and 5 min in 0.125 M glycine to sequence the reaction. Cells were lysed in 1% SDS, 10 mM EDTA, and 50 mM Tris-HCl (pH 8.0), and the DNA was fragmented to approximately 200-400 bp by sonication (Branson Sonifier 450). Immunoprecipitation was performed with 2 µg rabbit polyclonal anti-KDM2B (gift from Dr. Robert Klose, Oxford University) and rabbit anti-RING1B antibody (1:100, Cell Signaling Technology) overnight at 4°C. Antibody bound DNA-protein complexes were isolated by protein G plus/protein A agarose beads (EMD Millipore), washed and eluted, and reverse cross-linked DNA was extracted by phenol/chloroform and precipitated.

### ChIP DNA preparation for HiSeq4000 sequencing

ChIP DNA libraries were constructed for HiSeq4000 (Illumina) sequencing using NEBNext UltraII DNA library Prep Kit for Illumina (New England BioLabs, Inc) according to the manufacturer’s instructions. Adapter-ligated DNA was amplified by PCR for 12-14 cycles, followed by size selection using agarose gel electrophoresis. The DNA was purified using QIAquick gel extraction kits (Qiagen) and quantified both with an Agilent Bioanalyzer and Invitrogen Qubit. The DNA was diluted to a working concentration of 20nM prior to sequencing. Sequencing on an Illumina HiSeq4000 instrument was carried out by the Genomics Core Facility at Michigan State University.

### ChIP-Seq data analysis

For the ChIP-Seq data analysis, all sequencing reads were mapped to NCBI build 37 (mm9) of the mouse genome using Bowtie2^35^. Mapped reads were analyzed using the MACS program and bound-regions (peaks) were determined using sequencing reads from input DNA as negative controls^36^. When multiple reads mapped to the same genomic position, a maximum of two reads were retained. The statistical cutoff used for peak calling was *P*-value< 10^−8^ and >5-fold enrichment over the control. The mapped sequencing reads were normalized as Counts Per Million Reads (CPM). The normalized reads were binned into 50-bp windows along the genome using the bamCoverage of deepTools program and visualized in the IGV genome browser. The datasets of CpG islands and Refseq genes of mm9 mouse reference genome were retrieved from the UCSC table bowser. The heatmap and plot of ChIP-seq reads in the 10kb-windows flanking CGIs or Refseq genes were generated using plotHeatmap and plotProfile in the deepTools program^37^. The subset of bivalent promoters was determined by the 2kb promoter regions upstream of transcriptional start sites that contained both H3K27me3 and H3K4me3 using the bedtools program.

### RNA-seq sample preparation for HiSeq4000 sequencing

RNA was extracted and purified from cells using QI shredder (Qiagen) and RNeasy (Qiagen) spin columns. Total RNA (1 µg) was used to generate RNA-seq libraries using the NEBNext Ultra Directional RNA library Prep Kit for Illumina (New England BioLabs, Inc) according to the manufacturer’s instructions. Adapter-ligated cDNA was amplified by PCR and followed by size selection using agarose gel electrophoresis. The DNA was purified using the Qiaquick gel extraction kit (Qiagen) and quantified both with an Agilent Bioanalyzer and Invitrogen Qubit. The libraries were diluted to a working concentration of 10nM prior to sequencing. Sequencing on an Illumina HiSeq4000 instrument was carried out by the Genomics Core Facility at Michigan State University.

### RNA-Seq data analysis

RNA-Seq data analysis was performed essentially as described previously^38^. All sequencing reads were mapped mm9 of the mouse genome using Tophat2^39^. The mapped reads were normalized as Reads Per Kilobase of transcript per Million mapped reads (RPKM). The differential gene expression was calculated by the Cuffdiff program and the statistical cutoff for identification of differential gene expression was q < 0.05 and 2-fold RPKM change between samples^40^. The normalized mapped reads (RPKM) of each RNA-seq experiments were binned into 50bp windows along the genome using the bamCoverage of deepTools program and visualized in the IGV genome browser. The heatmap and plot of gene expression were generated using plotHeatmap and plotProile in the deepTools program^37^. The differentially expressed gene lists were then used as input for the DAVID Functional Annotation Bioinformatics Microarray Analysis for KEGG pathway and gene ontology enrichment analyses (https://david.ncifcrf.gov/).

## Acknowledgements

We thank MSU genomics core facility processed the next-generation sequencing. This work was supported by the National Institute Health NIH grant R01GM127431.

## Author contributions

J.H. conceived the project. The mouse model was generated in Y.Z. lab. N.D.W., and M.A. performed the experiments. J.H. and G.I.M. performed the sequencing data analysis. J.H., A.J.M., A.J.R., and Y.Z. discussed the results. Y.G. and J. H. wrote the manuscript.

