## Supplemental Fig.S1-S5 for "Impaired KDM2B-mediated PRC1 recruitment to chromatin causes neural stem cell senescence and ASD/ID-like behavioral deficits"

.

**This PDF file includes:**

Figs. S1 to S5
Table S1 to S5

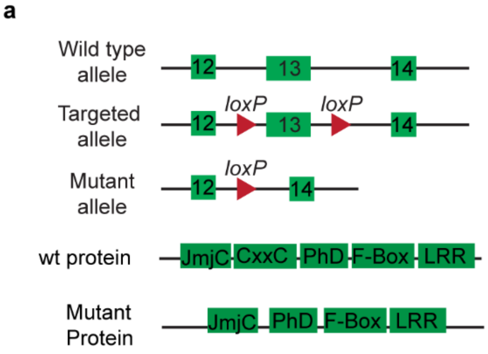


b

| Breeders | Nestin Cre+; Kdm2B fl/+ X Kdm2B fl/fl | | | |
|---|---|---|---|---|
| Genotype | Nestin Cre-; Kdm2B fl/+ | Nestin Cre-; Kdm2B fl/fl | Nestin Cre+; Kdm2B fl/+ | Nestin Cre+; Kdm2B fl/fl |
| Actual | 34 | 30 | 36 | 0 |
| Expect | 25 | 25 | 25 | 25 |

**Supplementary Figure 1. Characterization of *Kdm2b* conditional mutant mice.** (**a**) Diagram showing the strategy for the generation of *Kdm2b* conditional mutant mice. Cre-mediated deletion of exon 13 results in an altered spliced mRNA that encodes a mutant KDM2B protein with loss of CxxC-ZF domain. (**b**) The genotyping results of mice analyzed at postnatal stages.

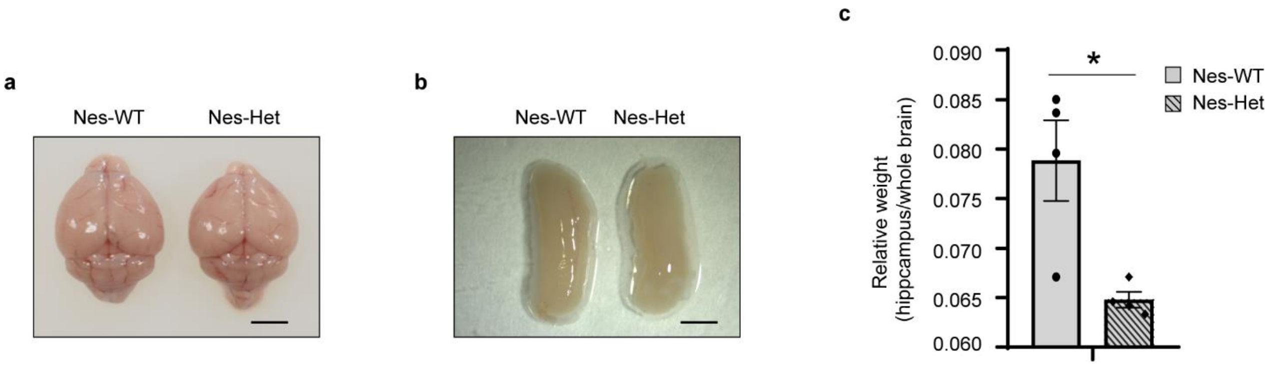


**Supplementary Figure 2. Loss of KDM2B CxxC-ZF domain induces cellular changes leading to NSC senescence.** (**a**) Photo showing the gross brain morphology of *Kdm2b*-Nes-WT and *Kdm2b*-Nes-Het mice at postnatal day 60, bar = 5 mm. (**b**) Photo showing the gross hippocampus morphology of *Kdm2b*-Nes-WT and *Kdm2b*-Nes-Het mice at postnatal day 60, bar = 1 mm. (c) Plot showing the relative weight of hippocampus normalized to whole brain. For each genotype, n = 4. *P*-values calculated using a two-tailed *t* test. Error bars in graphs represent mean ± SEM. Note: * $p < 0.05$.

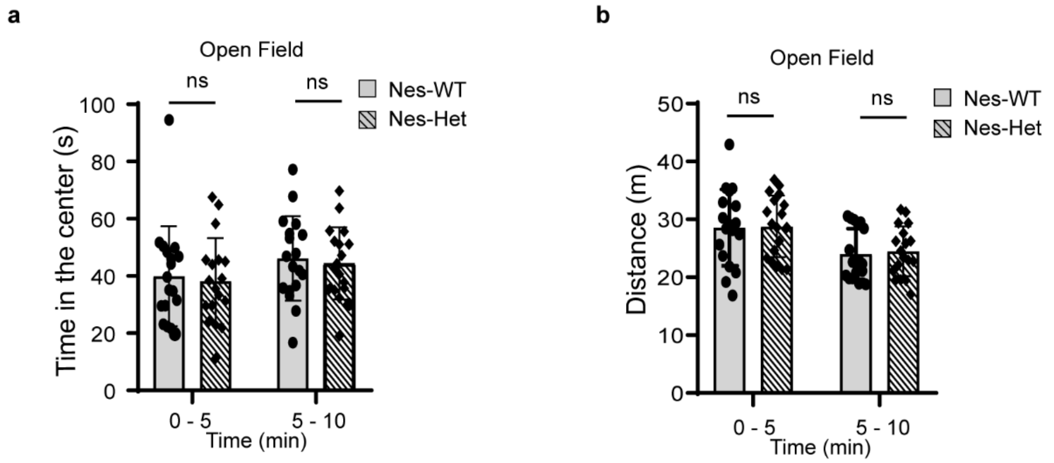


**Supplementary Figure 3. *Kdm*2b mutation in the developing brain causes ASD/ID-like behavioral and memory deficits.** (**a**) Time spent in the center of the open field arena measured in 10-min habituation. (**b**) Total distance traveled in 10-min habituation. For a and b, n = 15 for each genotype. *P*-values calculated using two-way ANOVA test. Error bars in graphs represent mean ± SEM. Note: ns, not significant.

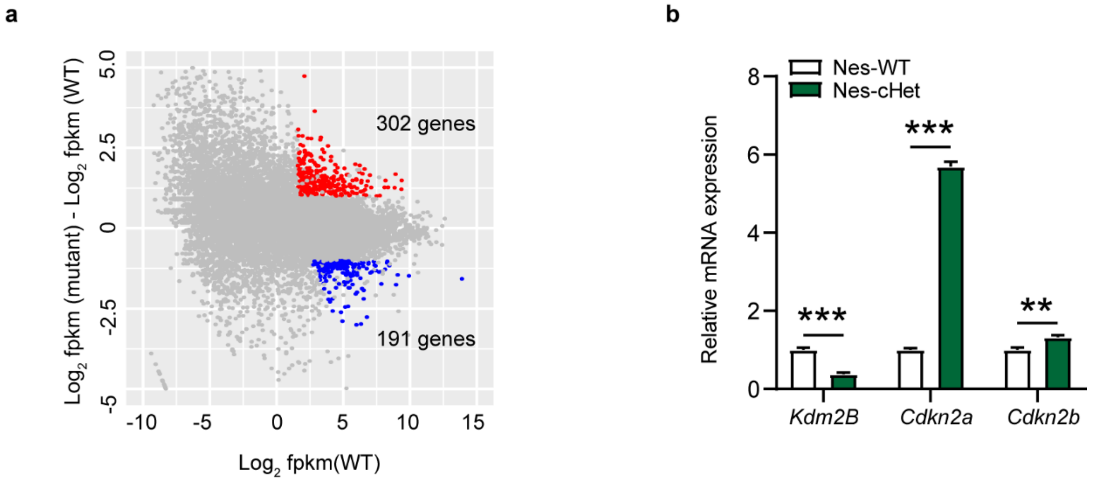


**Supplementary Figure 4. Loss of KDM2B CxxC-ZF domain depresses the genes involved in NSC senescence.** (**a**) Plot showing 302 up- and 191 down-regulated genes in the *Kdm2b*-Nes-Het NSCs compared to the wild-type NSCs. (**b**) qRT-PCR analysis showing the mRNA levels of *Kdm2b*, *Cdkn2a*, *and Cdkn2b* in *Kdm2b*-Nes-WT and *Kdm2b*-Nes-Het NSCs. The results of analysis in NSCs were normalized against levels of *Gapdh* and the expression level of wild-type NSCs was arbitrarily set to 1. The error bars represent mean ± SEM, n = 3 per genotype. *P*-values calculated using a two-tailed *t* test. ***P* < 0.01; ****P* < 0.001.

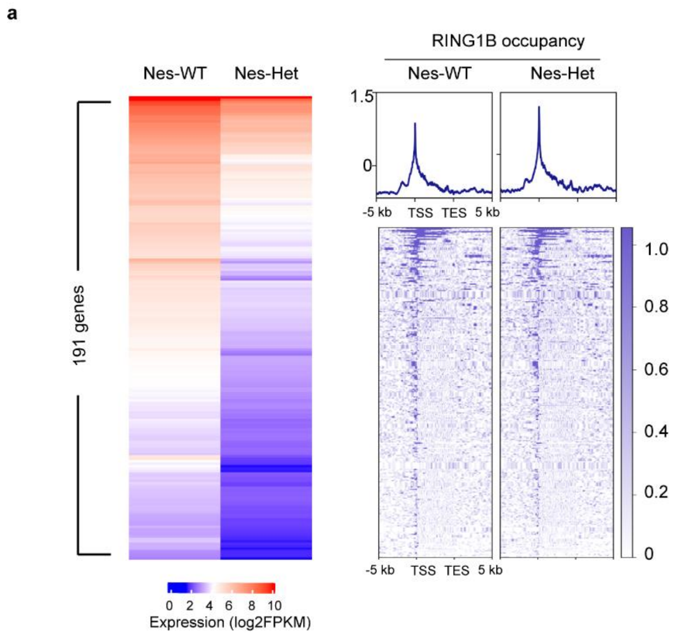


**Supplementary Figure 5. Loss of KDM2b CxxC-ZF domain impaired the PRC1 recruitment to chromatin. (a)** The heatmaps and plots showing the 191 downregulated genes in the *Kdm2b*-Nes-cKO NSCs (left panel) and the RING1B occupancy at the gene code regions and their 10-kb flanking regions (right panel). TSS: transcriptional starting sites; TES: transcriptional ending sites.

**Supplementary Table 1. Result of gene ontology enrichment analysis of genes upregulated in the *Kdm2b*-Nes-Het NSCs.**

| Term | Count | PValue | FDR |
|---|---|---|---|
| GO:0006915~apoptotic process | 27 | 5.34E-07 | 4.99E-04 |
| GO:0043065~positive regulation of apoptotic process | 18 | 1.37E-05 | 0.004276 |
| GO:0045669~positive regulation of osteoblast differentiation | 8 | 7.15E-05 | 0.014851 |
| GO:0008285~negative regulation of cell proliferation | 17 | 2.47E-04 | 0.034582 |
| GO:0051726~regulation of cell cycle | 9 | 2.92E-04 | 0.034582 |
| GO:0045597~positive regulation of cell differentiation | 6 | 6.15E-04 | 0.044381 |
| GO:0007399~nervous system development | 16 | 6.20E-04 | 0.044381 |

**Supplementary Table 2. Result of gene ontology enrichment analysis of genes down-regulated in the *Kdm2b*-Nes-Het NSCs.**

| Term | Count | PValue | FDR |
|---|---|---|---|
| GO:0007049~cell cycle | 24 | 1.30E-08 | 1.04E-05 |
| GO:0006260~DNA replication | 12 | 2.01E-08 | 1.04E-05 |
| GO:0007067~mitotic nuclear division | 12 | 5.77E-05 | 0.016998 |
| GO:0006270~DNA replication initiation | 5 | 6.61E-05 | 0.016998 |
| GO:0051301~cell division | 13 | 2.01E-04 | 0.041391 |

**Supplementary Table 3. Result of KEGG pathway enrichment analysis of genes upregulated in the *Kdm2b*-Nes-Het NSCs.**

| Term | Count | PValue | FDR |
|---|---|---|---|
| mmu04115:p53 signaling pathway | 8 | 2.49E-05 | 0.004338 |
| mmu04068:FoxO signaling pathway | 10 | 6.09E-05 | 0.005296 |
| mmu04010:MAPK signaling pathway | 12 | 4.25E-04 | 0.018486 |
| mmu04024:cAMP signaling pathway | 10 | 0.00108 | 0.037587 |
| mmu04151:PI3K-Akt signaling pathway | 13 | 0.002062 | 0.039874 |

**Supplementary Table 4. Result of KEGG pathway enrichment analysis of genes down-regulated in the *Kdm2b*-Nes-Het NSCs.**

| Term | Count | PValue | FDR |
|---|---|---|---|
| mmu03030:DNA replication | 5 | 2.52E-04 | 0.035772 |
| mmu00100:Steroid biosynthesis | 4 | 6.04E-04 | 0.042901 |

**Supplementary Table 5. Sequences of all primers used in this study.**

| Name | Sequence (5'-3') | Purpose |
|---|---|---|
| GAPDH-F | gcagtggcaaagtggagatt | qRT-PCR |
| GAPDH-R | gaatttgccgtgagtggagt | qRT-PCR |
| Cdkn2a-F | gaactctttcggtcgtaccc | qRT-PCR |
| Cdkn2a-R | cgaatctgcaccgtagttga | qRT-PCR |
| Cdkn2b-F | ccacccttaccagacctgtg | qRT-PCR |
| Cdkn2b-R | aggcgtcacacacatccag | qRT-PCR |
| Nes-cre-F | gcggtctggcagtaaaaactatc | Genotyping |
| Nes-cre-R | gtgaaacagcattgctgtcactt | Genotyping |
| Kdm2b-mut-F | taactgtggcaggaggaggt | Genotyping |
| Kdm2b-mut_R | caacttgaacctttggagca | Genotyping |
